# 5-methylcytosine modification by *Plasmodium* NSUN2 stabilizes mRNA and mediates the development of gametocytes

**DOI:** 10.1101/2021.06.06.447275

**Authors:** Meng Liu, Gangqiang Guo, Pengge Qian, Jianbing Mu, Binbin Lu, Xiaoqin He, Xiaomin Shang, Guang Yang, Shijun Shen, Wenju Liu, Liping Wang, Liang Gu, Quankai Mu, Xinyu Yu, Yuemeng Zhao, Richard Culleton, Jun Cao, Lubin Jiang, Thomas E. Wellems, Jing Yuan, Cizhong Jiang, Qingfeng Zhang

**Affiliations:** Key Laboratory of Spine and Spinal Cord Injury Repair and Regeneration of Ministry of Education, Orthopaedic Department of Tongji Hospital, Shanghai Key Laboratory of Signaling and Disease Research, School of Life Sciences and Technology, Tongji University, Shanghai 200065, China; Unit of Molecular Parasitology, Research Center for Translational Medicine, East Hospital, Tongji University School of Medicine, Shanghai 200120, China; State Key Laboratory of Cellular Stress Biology, Innovation Center for Cell Signaling Network, School of Life Sciences, Xiamen University, Xiamen, Fujian, China; Laboratory of Malaria and Vector Research, National Institute of Allergy and Infectious Diseases, National Institutes of Health, Rockville, MD 20892-8132, USA; National Health Commission Key Laboratory of Parasitic Disease Control and Prevention, Jiangsu Provincial Key Laboratory on Parasite and Vector Control Technology, Jiangsu Institute of Parasitic Diseases, Wuxi, 214064, China; Department of Immunogenetics, Institute of Tropical Medicine (NEKKEN), Nagasaki University, 1-12-4 Sakamoto, Nagasaki 852-8523, Japan; Unit of Human Parasite Molecular and Cell Biology, Key Laboratory of Molecular Virology and Immunology, Institut Pasteur of Shanghai, University of Chinese Academy of Sciences, Chinese Academy of Sciences, Shanghai 200031, China; Division of Molecular Parasitology, Proteo-Science Centre, Ehime University, Matsuyama, Ehime 790-8577, Japan; Department of Protozoology, Institute of Tropical Medicine, Nagasaki University, Nagasaki, Japan

**Author notes:** These authors contributed equally to this work (Meng Liu, Gangqiang Guo, Pengge Qian, Jianbing Mu).

**Keywords:** RNA methyltransferase, gene knock-out, gametocytogenesis, epitranscriptomic modifications, m^5^C transcript profiles, RNA bisulfite sequencing

## Abstract

5-methylcytosine (m^5^C) is an important epitranscriptomic modification involved in mRNA stability and translation efficiency in various biological processes. However, it remains unclear if m^5^C modification contributes to the dynamic regulation of the transcriptome during the developmental cycles of *Plasmodium* parasites. Here, we characterize the landscape of m^5^C mRNA modifications at single nucleotide resolution in the asexual replication stages and gametocyte sexual stages of rodent (*P. yoelii*) and human (*P. falciparum*) malaria parasites. While different representations of m^5^C-modified mRNAs are associated with the different stages, the abundance of the m^5^C marker is strikingly enhanced in the transcriptomes of gametocytes. Our results show that m^5^C modifications confer stability to the *Plasmodium* transcripts and that a *Plasmodium* ortholog of NSUN2 is a major mRNA m^5^C methyltransferase in malaria parasites. Upon knock-out of *P. yoelii nsun2* (*pynsun2*), marked reductions of m^5^C modification were observed in a panel of gametocytogenesis-associated transcripts. These reductions correlated with impaired gametocyte production in rodent and human malaria parasites. Restoration of the *nsun2* gene in the knock-out parasites rescued the gametocyte production phenotype as well as m^5^C modification of the gametocytogenesis-associated transcripts. Together with the mRNA m^5^C profiles for two species of *Plasmodium*, our findings demonstrate a major role for NSUN2-mediated m^5^C modifications in mRNA transcript stability and sexual differentiation in malaria parasites.

**Significance:** Modifications of RNA including methylations of cytosine (m^5^C) and adenosine (m^6^A) have important roles in RNA metabolism, cellular responses to stress, and biological processes of differentiation and development. Here, we report on the profiles of m^5^C mRNA modifications in malaria parasites that infect rodents (*Plasmodium yoelii*) and humans (*Plasmodium falciparum*). These parasites have genes that encode homologs of human and plant NSUN2 methyltransferases (m^5^C “writers”). We show that one of these homologs, termed PyNSUN2, stabilizes mRNA transcripts in *P. yoelii* and mediates m^5^C-associated development of the parasite sexual stages (gametocytes). Further research on m^5^C and other epitranscriptomic modifications may yield new insights into molecular pathways of gametocyte development and mosquito infectivity that can be exploited to interrupt malaria transmission.

## Introduction

Malaria is a mosquito-borne infectious disease caused by unicellular apicomplexan protozoa of the genus *Plasmodium*. Consequences of this disease in 2019 included an estimated 229 million infections and 409,000 deaths globally (1). Among the species infecting humans, *Plasmodium falciparum* is the most virulent and accounts for most of these deaths. *Plasmodium* parasites have a complex life cycle that alternates between mosquito and vertebrate hosts, in which the parasites complete numerous rounds of asexual proliferation in the haploid state and pass through a brief diploid period following obligatory mating of male and female gametes in the mosquito midgut (2). During the proliferation of intraerythrocytic parasites in the vertebrate bloodstream, a small fraction of the asexual population commits to gametocytogenesis, producing sexual stage gametocytes that are taken up by mosquitos in which they emerge from red blood cells as gametes (3, 4). These cellular developments are associated with stage-specific and highly dynamic transcriptomes under the hierarchical control of transcription factors and epigenetic regulators (3, 5, 6).

Epigenetic regulation at the transcriptional level plays a critical role in genome expression events and their global outcome. For instance, heterochromatin protein 1 (HP1)-dependent heterochromatin modification provides a transcriptionally repressive microenvironment for the silencing of antigenically variant genes and *ap2-g*, a master regulator of sexual commitment, thus determining parasite adaptation and development in the human host and transmission into mosquitoes (5, 7, 8). Regulatory pathways such as RNase-mediated nascent decay, m6A modification-mediated mRNA stability, and RNA helicase-associated translational repression confer other important mechanisms for fine-tuning post-transcriptional regulation in malaria parasites (9–11). These findings demonstrate that the highly dynamic transcriptome of malaria parasites is controlled by a complex but well-organized multi-layered regulatory network.

While DNA methylation has been widely studied as an epigenetic phenomenon, characterizations of mRNA methylation in the modulation of transcriptome stability and expression are more recent (12–14). Reversible mRNA modifications are now known to be involved in the post-transcriptional regulation of gene expression in eukaryotes (11). For example, N6-methyladenosine (m^6^A) is the most prevalent RNA modification and has been investigated extensively in many eukaryotic organisms, particularly in its association with various cellular processes such as cell differentiation, embryonic development, and stress responses. In these processes m6A epitranscriptomic modifications are involved in regulation of mRNA metabolism, translation, decay, and miRNA biogenesis (11, 15, 16). In malaria parasites, the epitranscriptome has likewise been recognized as an important modulator of post-transcriptional gene expression. Baumgarten *et al.* (12) described highly regulated features of m^6^A mRNA methylation in *P. falciparum*, found that these features are associated with stage-specific fine-tuning of the transcriptional cascade, and suggested that widely distributed m6A modifications may shape transcriptome expression through effects on mRNA stability during blood-stage development.

In addition to m^6^A, RNA modifications such as N1-methyladenosine (m^1^A), m^5^C, 5-hydroxymethylcytosine (hm^5^C), pseudouridine (Ψ), inosine (I), uridine (U) and methylation of ribose at the 2’ position (2’-O-Me) are also known to regulate transcription (15). Modification of mRNA by m^5^C is important for diverse biological processes including the progression of bladder cancer, HIV-1 infection, and developmental progressions in zebrafish, *Arabidopsis thaliana*, and rice (13, 17–19), suggesting that cytosine methylation is a powerful regulator of cellular processes at the epitranscriptomic level (16). In other studies, pathways involving m^5^C-modified transcripts have been found to include DNA repair and homologous recombination (20). Specific methyltransferases (m^5^C writers) in various species include NSUN2 in humans, TRM48 in *Arabidopsis thaliana*, and OsNSUN2 in rice. NSUN2 can drive human urothelial carcinoma pathogenesis by targeting the m^5^C methylation site in the HDGF 3′ untranslated region (18). TRM4B, by promoting mRNA stability through m^5^C modifications, influences the transcriptional levels of genes involved in the root development of *Arabidopsis thaliana* (13). Mutation of OsNSUN2 affects photosynthesis efficiency in rice (19).

Among binding proteins (m^5^C “readers”) that recognize m^5^C-modified transcripts are the Aly/REF export factor (ALYREF) in humans and the Y-box binding protein 1 (Ybx1) in zebrafish. ALYREF is an adaptor that regulates the transport of m^5^C-modified transcripts from the nucleus to the cytoplasm (14), whereas Ybx1 promotes the stability of its target mRNAs in an m^5^C-dependent manner (21). A potential m^5^C demethylase, ALKBH (an AlkB homolog), was reported more recently from *Arabidopsis* and is attracting study for its possible role as an m^5^C “eraser” (22, 23).

In *P. falciparum*, the DNA methyltransferase homologue TRDMT1 has been shown to modify position 38 of aspartic acid tRNA, where the presence of m^5^C may modulate translation of proteins with effects on infection and pathogenesis (24). More generally, a number of RNA (cytosine-5)-methyltransferase (RCMT) orthologs have been predicted from genome analysis of *Plasmodium* species including *P. falciparum* and the rodent malaria parasite *Plasmodium yoelii* (25–27). Findings from these studies suggest that m^5^C mRNA methylations occur, but the patterns and roles of m^5^C modifications in the biological processes of parasite infection, virulence, and transmission remain largely unknown. Among *Plasmodium* species, *P. falciparum* and *P. yoelii* have comparatively low G+C contents (both approximately 20%) (25). Considering these low G+C contents, it is interesting to speculate whether epitranscriptomic effects from m^5^C modifications may differ in these parasites compared to those in other *Plasmodium* species that have higher G+C contents.

The present study was designed to characterize the overall transcriptome profile and stage-specific dynamics of m^5^C mRNA modifications in the asexual replicating forms and gametocytes of *P. yoelii* and *P. falciparum*. Our results demonstrate comparatively high levels of m^5^C mRNA methylation in the gametocytes of both species. Enhanced stability of m^5^C transcripts is likely to play an important role in the processes of sexual stage development and transmission. *P. yoelii* and *P. falciparum* each carry an ortholog of NSUN2 (PyNSUN2; PfNSUN2) that functions as a methyltransferase with a major role in these m^5^C modifications of the transcriptome. We show that *P. yoelii* gametocytogenesis is disrupted by knock-out of the *Pynsun2* gene and can be reestablished by restoration of *Pynsun2* expression.

## Results

### Global profiles of m^5^C modification in the *P. yoelii* and *P. falciparum* transcriptomes

To test for detectable m^5^C modifications, dot blot assays were performed with mRNA preparations from *P. yoelii* and *P. falciparum* mRNAs. Using an anti-m^5^C antibody previously shown to detect mRNA m^5^C modifications in *Oryza sativa* (19), our experiments demonstrated the presence of m^5^C modifications in schizont-stage parasites of both *Plasmodium* species (Fig. 1*A*).

**Figure 1.**
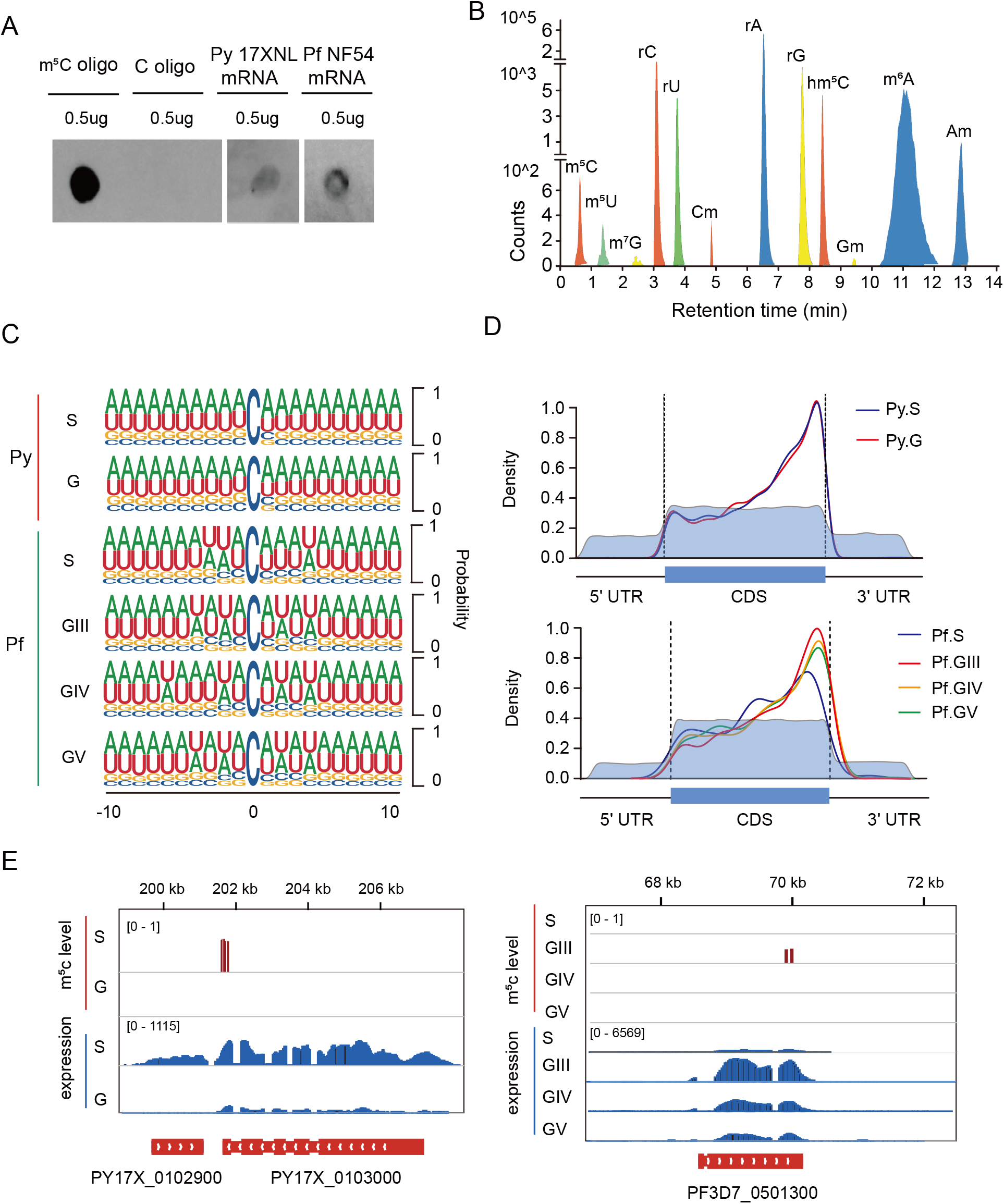
Features of m^5^C-modified mRNA in malaria parasites. **A**, Dot blot assays show detection of m^5^C modifications in schizont-stage mRNAs from *P. falciparum* and *P. yoelii*. m^5^C-modified and unmodified RNA oligonucleotides served as positive and negative controls, respectively. **B**, LC-MS/MS signals indicating modified nucleotides from *P. yoelii* schizont-stage mRNA. rC, cytosine; rU, uracil; rG, guanosine; rA, adenosine; m⁶A, N⁶-methyladenosine; m⁷G, N⁷-methyl-2’-guanosine; Am, 2’-O-methyladenosine; m⁵C, 5-methylcytosine; Gm, 2’-O-methylguanosine; m⁵U, 5-methyluridine; Cm, 2’-O-methylcytosine; hm^5^C, 5-Hydroxymethylcytosine. **C**, Frequency logo displays of the nucleotides proximal to mRNA m^5^C sites in the schizonts (S) and gametocytes (G) of *P. yoelii* (Py), and the schizonts (S) and stage III, IV, and V gametocytes (GIII, GIV, GV) of *P. falciparum* (Pf). **D**, Density distribution of the m^5^C sites in mRNA transcripts of *P. yoelii* and *P. falciparum*. The moving averages (10-bp window) of percentage mRNA cytosine content (light blue) are lower in the 5’ and 3’ UTR regions than in the CDS, as expected for these species. **E**, Integrative Genomics Viewer snapshots of the transcript sequence levels and m^5^C sites of expressed genes in *P. yoelii* (left) and *P. falciparum* (right). Vertical red bars indicate the m^5^C levels in specific parasite stages.

For quantitative assessments of modified nucleotides in the *P. yoelii* transcriptome, purified mRNA preparations from *P. yoelii* were completely digested to mononucleotides, which were then dephosphorylated and subjected to analysis by triple quadrupole liquid chromatography–mass spectrometry (QQQ LC–MS). Results identified well-known mRNA modifications including m^6^A, the most abundant mRNA modification in eukaryotes including *P. falciparum* (12, 28). Other modifications including m^5^C, hm^5^C, m^5^U, and m^7^G were detected at lower levels (Fig. 1*B*).

To obtain a transcriptome-wide maps of m^5^C modification at single-base resolution, we performed RNA-BisSeq on mRNA preparations from asexual schizont (S) and sexual gametocyte (G) stages of *P. yoelii*, and from synchronized schizonts and three stages of induced gametocytes (5–6 day stage III, 8–9 day stage IV, and 12–13 day stage V) of *P. falciparum*. For each stage, two biological replicates were used for high-confidence site calling (SI Appendix, Table S1). The m^5^C sites identified between the independent replicates displayed variation and reproducibility comparable to those of high quality studies from other organisms such as zebrafish, human, and mouse (21, 29). About one third of m^5^C sites were shared between replicates (SI Appendix, Fig. *S1A*). The methylation levels of these shared (common) m^5^C sites were significantly higher than those of the m^5^C sites unique to one replicate or the other (SI Appendix, Fig. *S1A*, box plots). Median fractional methylation levels at the shared sites were 0.77 for *P. yoelii* schizonts and 0.15 for *P. falciparum* schizonts. In comparison, the median m^5^C methylation levels were 0.39 for *P. yoelii* day 3 gametocytes, and 0.2, 0.48, and 0.39 for the stage III, IV, and V *P. falciparum* gametocytes, respectively (SI Appendix, Fig. *S1B*).

Examination of complete transcripts harboring m^5^C modifications showed that individual transcripts more often carried these modifications at two or more sites than at just one site alone, and that a greater fraction of *P. yoelii* transcripts carried multiple m^5^C modifications than did *P. falciparum* transcripts (SI Appendix, Fig. S1*C* and SI Appendix, Table S1). Frequency analyses of the nucleotides neighboring the modifications found that most m^5^C sites were in CHH regions (where H = A, C, U) (SI Appendix, Fig. S1*D*) and often in the context of AU-rich segments (Fig. 1*C*). The large majority of modifications were identified within the transcript coding sequences (CDSs; SI Appendix, Fig. S1*E*), where most of these m^5^C sites were located upstream of stop codons (Fig. 1*D*) in a different pattern than in human HeLa cells (14) and zebrafish embryos (21). Fig. 1E presents representative examples of *P. yoelii* and *P. falciparum* transcripts that carry m^5^C modifications in the 3’ regions of their CDSs.

### Profiles of m^5^C modifications in the asexual and sexual stages of *P. yoelii* and *P. falciparum*

Results of RNA-BisSeq analysis identified 7409 m^5^C modifications in 527 mRNA transcripts from the *P. yoelii* schizont set (Py17X_S; SI Appendix, Table S2) and 9168 m^5^C modification in 796 transcripts from the *P. yoelii* gametocyte set (Py17X_G; SI Appendix, Table S2). In contrast, results from the *P. falciparum* datasets showed lower levels of 335 m^5^C modifications in 220 asexual stage transcripts, and 781, 419, and 891 m^5^C modifications in 422 GIII, 224 GIV, and 438 GV transcripts, respectively (SI Appendix, Table S2).

To further investigate potential roles of mRNA m^5^C modifications in the *Plasmodium* life cycle, we performed Gene Ontology (GO) analysis on the sets of methylated transcripts from *P. yoelii* and *P. falciparum.* For *P. yoelii* gametocytes, results from this analysis identified enriched numbers of transcripts relating to metabolic/cellular processes and sexual development. Correspondingly, for *P. falciparum* gametocytes, transcripts relating to cellular and metabolic processes were enriched in GIII stages and transcripts relating to sexual development were enriched in GV stages (Table 1). In the asexual stage parasites including schizonts, m^5^C modifications were enriched with transcripts having assigned GO terms of nucleoside transport, nucleic acid metabolic processes, and chromatin organization (Table 1). These features were also evident in heatmaps of the m^5^C-modified transcripts, which identified similarities as well as distinct differences between the asexual stages and gametocytes both within and between the *P. yoelii* and *P. falciparum* populations (SI Appendix, Fig. S2).

**Table 1.**
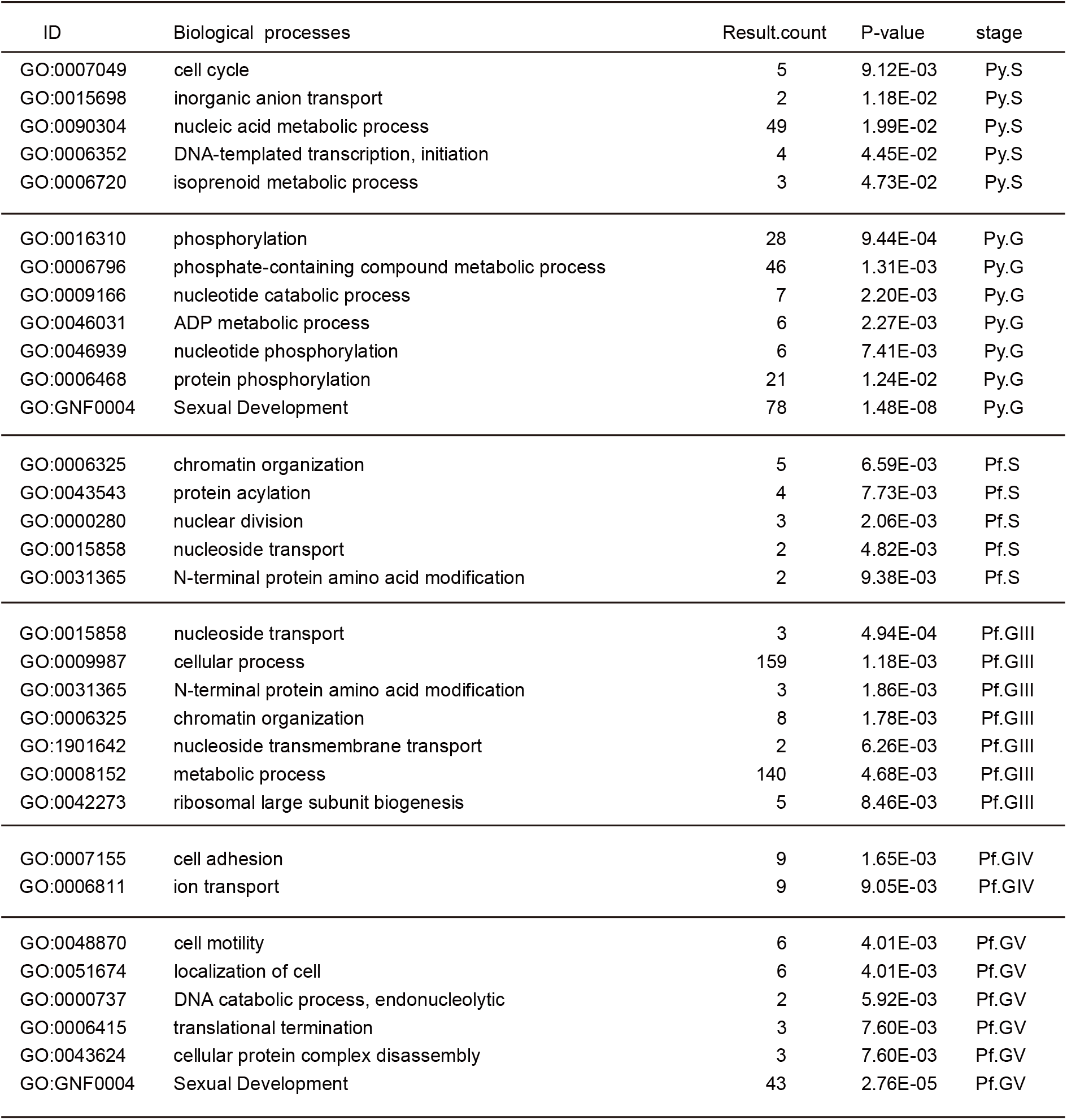
Gene Ontology (GO) for m^5^C-modified transcripts in schizonts and gametocytes of *P. yoelii* (E) and *P. falciparum* (F).

| ID | Biological processes | Result.count | P-value | stage |
| --- | --- | --- | --- | --- |
| GO:0007049 | cell cycle | 5 | 9.12E-03 | Py.S |
| GO:0015698 | inorganic anion transport | 2 | 1.18E-02 | Py.S |
| GO:0090304 | nucleic acid metabolic process | 49 | 1.99E-02 | Py.S |
| GO:0006352 | DNA-templated transcription, initiation | 4 | 4.45E-02 | Py.S |
| GO:0006720 | isoprenoid metabolic process | 3 | 4.73E-02 | Py.S |
| GO:0016310 | phosphorylation | 28 | 9.44E-04 | Py.G |
| GO:0006796 | phosphate-containing compound metabolic process | 46 | 1.31E-03 | Py.G |
| GO:0009166 | nucleotide catabolic process | 7 | 2.20E-03 | Py.G |
| GO:0046031 | ADP metabolic process | 6 | 2.27E-03 | Py.G |
| GO:0046939 | nucleotide phosphorylation | 6 | 7.41E-03 | Py.G |
| GO:0006468 | protein phosphorylation | 21 | 1.24E-02 | Py.G |
| GO:GNF0004 | Sexual Development | 78 | 1.48E-08 | Py.G |
| GO:0006325 | chromatin organization | 5 | 6.59E-03 | Pf.S |
| GO:0043543 | protein acylation | 4 | 7.73E-03 | Pf.S |
| GO:0000280 | nuclear division | 3 | 2.06E-03 | Pf.S |
| GO:0015858 | nucleoside transport | 2 | 4.82E-03 | Pf.S |
| GO:0031365 | N-terminal protein amino acid modification | 2 | 9.38E-03 | Pf.S |
| GO:0015858 | nucleoside transport | 3 | 4.94E-04 | Pf.GIII |
| GO:0009987 | cellular process | 159 | 1.18E-03 | Pf.GIII |
| GO:0031365 | N-terminal protein amino acid modification | 3 | 1.86E-03 | Pf.GIII |
| GO:0006325 | chromatin organization | 8 | 1.78E-03 | Pf.GIII |
| GO:1901642 | nucleoside transmembrane transport | 2 | 6.26E-03 | Pf.GIII |
| GO:0008152 | metabolic process | 140 | 4.68E-03 | Pf.GIII |
| GO:0042273 | ribosomal large subunit biogenesis | 5 | 8.46E-03 | Pf.GIII |
| GO:0007155 | cell adhesion | 9 | 1.65E-03 | Pf.GIV |
| GO:0006811 | ion transport | 9 | 9.05E-03 | Pf.GIV |
| GO:0048870 | cell motility | 6 | 4.01E-03 | Pf.GV |
| GO:0051674 | localization of cell | 6 | 4.01E-03 | Pf.GV |
| GO:0000737 | DNA catabolic process, endonucleolytic | 2 | 5.92E-03 | Pf.GV |
| GO:0006415 | translational termination | 3 | 7.60E-03 | Pf.GV |
| GO:0043624 | cellular protein complex disassembly | 3 | 7.60E-03 | Pf.GV |
| GO:GNF0004 | Sexual Development | 43 | 2.76E-05 | Pf.GV |

### m^5^C-mediated mRNA stability correlates with gametocyte development

Having found that m^5^C methylated transcripts were significantly enriched for GO identifications of sexual development in *P. yoelii* and *P. falciparum* gametocytes, we focused next on genes previously reported to be involved in gametocyte development (30). Our search of the *P. yoelii* data identified 848 genes with transcript abundances of more than 3-fold (log_2_ ≥1.585) in gametocytes relative to asexual parasites (SI Appendix, Table S3). Similarly, 661 gametocyte development-related genes were identified with ≥ 3-fold higher transcript abundance in *P. falciparum* gametocytes (stage GIII) *vs.* asexual stages (SI Appendix, Table S3). On comparative analysis of these gametocyte transcripts, significantly increased m^5^C levels were observed relative to the schizont transcripts of *P. yoelii* and *P. falciparum*, particularly in the comparison using m^5^C-modified stage V gametocyte transcripts (Fig. 2*A* and *B*).

**Figure 2.**
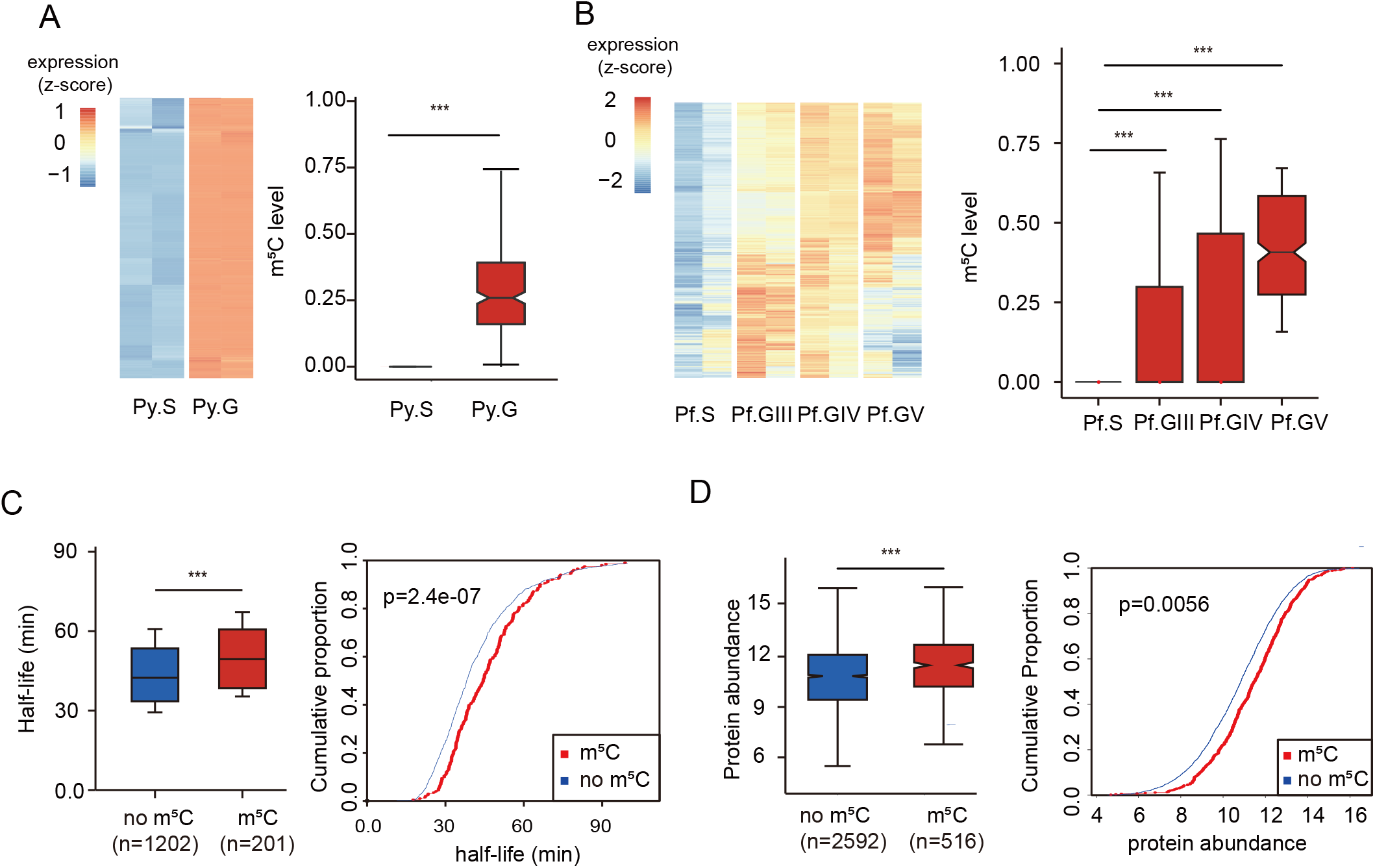
High levels of m^5^C methylation and its association with transcript longevity and protein expression in gametocytes. **A-B**, Heatmaps indicate the expression of transcripts whose levels are ≥ 3-fold higher in gametocytes than in schizonts of *P. yoelii* and *P. falciparum.* Boxplots show corresponding m^5^C levels of those genes at different stages. (***: p-value < 0.001, Wilcoxon test). **C**, Boxplot and cumulative fraction plot indicate the longer mRNA half-lives of m^5^C methylated (red) relative to non-methylated (blue) transcripts in *P. yoelii* gametocytes (***: p-value < 0.001, Wilcoxon test.). **D**, Boxplot and cumulative fraction plot compare the levels of proteins translated from m^5^C-methylated (red) and non-methylated (blue) transcripts in *P. yoelii* gametocytes (***: *P* < 0.001, Wilcoxon test).

Our comparisons confirmed the presence of m^5^C modifications in transcripts with well-established roles in sexual commitment (3, 5, 6, 31). For example, m^5^C methylation of AP2-O (PY17X_0907300) transcripts was not detected in schizonts but was present at a level of 0.34 along with higher expression of these transcripts in gametocytes (SI Appendix, Fig. S3*A*). In another example, the DEAD/DEAH helicase (PY17X_0313200), an AP2-G-induced transcript involved in male gametocyte development in *P. berghei* (5), was found to be m^5^C methylated in both asexual and sexual stages of *P. yoelii*. Similarly, data from *P. falciparum* showed stage-specific m^5^C methylation of three transcripts that possibly regulate sexual commitment and gametocytogenesis (2, 3, 23), including genes whose disruption is known to lead to greatly reduced gametocyte numbers (GIG, AP2-G2, Pfs16) (2, 3) (SI Appendix, Fig. S3*A*). However, no detectable m^5^C modification was found for AP2-G transcripts in either the *P. yoelii* or *P. falciparum* malaria parasites.

In view of the above results and m^5^C stabilization of transcripts known from other systems (18, 21), we asked if m^5^C might be associated with mRNA stability in *Plasmodium* parasites. To address this question, we first assessed the abundance of mRNAs whose m^5^C levels differed in sexual and asexual parasites, relative to the abundance of mRNAs that showed no change of m^5^C levels between these stages. Results from these comparisons suggested a positive correlation between m^5^C level and mRNA abundance in malaria gametocytes relative to schizonts (SI Appendix, Fig. S3*B*).

We next studied the half-lives of mRNAs in *P. yoelii* gametocytes that were treated with actinomycin D as a transcription inhibitor (18). Results from RNA-Seq analysis indicated that mRNAs with m^5^C modification had significantly longer half-lives than those without m^5^C modification (Fig. 2*C* and SI Appendix, Table S4). Protein quantification with Tandem Mass Tags (TMT) (SI Appendix, Table S5) showed that slightly greater protein abundance in *P. yoelii* was associated with m^5^C-modified than unmodified transcripts (Fig. 2*D*).

### PyNSUN2 functions as a mRNA m^5^C methyltransferase in *P. yoelii*

To identify potential mRNA m^5^C methyltransferases in *P. yoelii*, we searched for orthologs of hNSUN2 and identified four candidate sequences: PY17X_0804600, PY17X_0938600, PY17X_0920500, PY17X_1447800 (named PyNSUN1–PyNSUN4, respectively) (Fig. 3*A*). Corresponding searches of the www.plasmoDB.org database identified PF3D7_0704200, PF3D7_1111000, PF3D7_1129400, and PF3D7_1230600 (PfNSUN1–PfNSUN4) as candidate RNA methyltransferases in *P. falciparum*. Sequence comparisons showed that individual members of this family are highly conserved between *P. yoelii* and *P. falciparum*, and that the NSUN1 and NSUN2 sequences have close affinities to orthologs in humans, *Arabidopsis thaliana*, *Saccharomyces cerevisiae*, and *Oryza sativa* (Fig. 3*A*).

**Figure 3.**
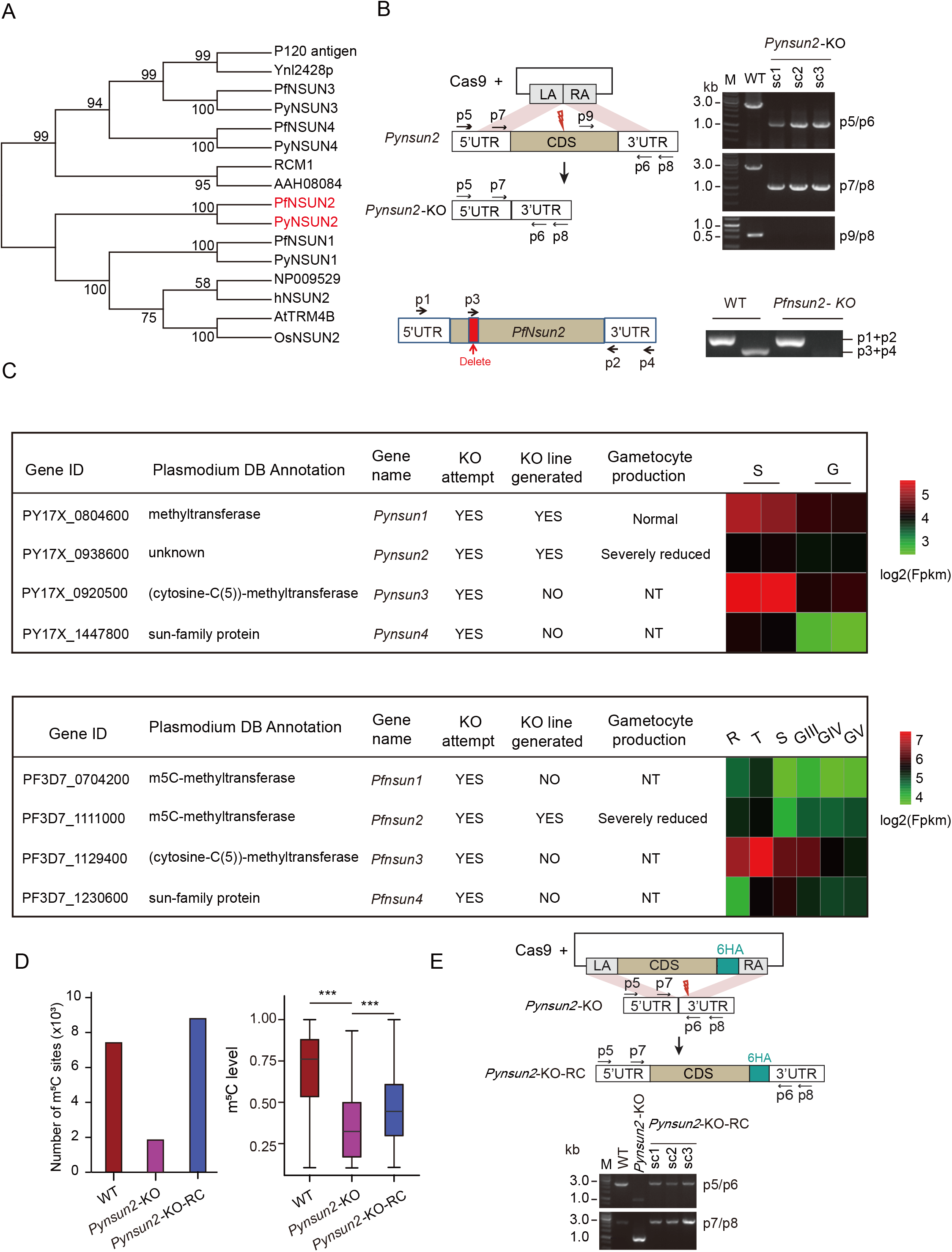
Identification and verification of PyNSUN2 as a m^5^C methyltransferase. **A**, Phylogenetic analysis of putative RNA (cytosine-5)-Methyltransferases (NSUNs) of *P. yoelii* and *P. falciparum*. Sequences were aligned using ClustalX 2.1. The neighbor joining (NJ) phylogeny was performed using MEGA 5.2.2 with 1000 replicates. **B**, Strategies to generate *Pynsun2*-KO and *Pfnsun2*-KO parasites. For CRISPR/Cas9-mediated deletion of the *Pynsun2* CDS, Left Arm (LA) and Right Arm (RA) match sequences in the 5’ UTR and 3’ UTR of *Pynsun2*, respectively. For disruption of *Pfnsun2*, sgRNA targeting was used to delete a portion of the CDS (red box) and introduce a stop codon. PCR products with indicated primer pairs confirmed the expected differences between the WT and allelically-manipulated parasites. Sequence data confirming the deletion and stop codon in the *Pfnsun2-*KO parasites is shown in SI Appendix, Fig. S4*A.* **C**, Summary results from experiments to disrupt homologs of the NSUN family in *P. yoelii* and *P. falciparum.* NT: not tested. **D**, Histogram shows numbers of mRNA m^5^C sites in schizont stages of WT, *Pynsun2*-KO and *Pynsun2*-KO-RC *P. yoelii clones*. Box plot shows the corresponding m^5^C levels in WT, *Pynsun2*-KO and *Pynsun2*-KO-RC lines (right panel). (***: *P*-value < 0.001, Wilcoxon test.) **E**, Strategy for genetic complementation repair of the *Pynsun2-*KO with the gene CDS fused with a C-terminal 6HA. Upper panel shows the design for the CRISPR/Cas9 mediated gene knock-in. Left Arm (LA) and Right Arm (RA) match sequences in the 5’ UTR and 3’ UTR of *Pynsun2*, respectively. Red thunderbolt indicates the site for sgRNA targeting. Bottom panel shows the PCR products in the three complemented clones (RC) by the indicated primer sets.

To study the role of these putative m^5^C methyltransferases in *Plasmodium*, we performed experiments to disrupt *Pynsun2* in the *P. yoelii* 17XNL line and *Pfnsun2* in the *P. falciparum* NF54 line by CRISPR-Cas9 gene editing (Fig. 3*B*). Functional disruption attempts of the *sun1*, *sun3*, and *sun4* homologs were performed similarly. After a minimum of three independent transfection attempts with 2 to 3 different single guide RNA (sgRNA) sequences for each individual homolog (SI Appendix, Table S6), we obtained knock-out lines for three of the eight genes (*Pynsun1*, *Pynsun2* and *Pfnsun2*) (Fig. 3*C* and SI Appendix, Fig. S4*A*).

Comparative RNA-BisSeq analysis on the schizont stages of 17XNL wild-type (WT) and *Pynsun2* knock-out parasites demonstrated a marked decrease in the density of m^5^C modifications in mRNA transcripts (Fig. 3*D*). By RNA-BisSeq, 7409 m^5^C sites were detected in 527 mRNAs of the WT parasites whereas this number was reduced to 1845 m^5^C sites in 265 mRNAs of the knock-outs (SI Appendix, Table S2).

To confirm the evidence from knock-out parasites that *Pynsun2* functions as a major m^5^C methyltransferase, we re-introduced *Pynsun2* to its original locus using CRISPR-Cas9 allelic modification (Fig. 3*E* and SI Appendix, Fig. S4*B*). Comparative analysis of the restored line (*Pynsun2-*KO-RC) confirmed the return of substantial m^5^C levels, reflecting restored methylation activity (Fig. 3*D*).

### Gametocytogenesis is impaired by disruption of *Plasmodium nsun2* and can be restored by genetic complementation

Considering the differentially greater m^5^C levels in gametocyte *vs.* schizont mRNA transcripts and the successful knock-outs of *Pynsun1*, *Pfnsun2*, and *Pfnsun2* in blood stage parasites, we asked if expression of these genes might be associated with phenotypes of gametocyte production or parasite stages development in the mosquito. Comparative experiments with the knock-out and WT lines demonstrated that disruptions of *Pynsun2* in *P. yoelii* and *Pfnsun2* in *P. falciparum* resulted in dramatically reduced gametocyte production (Fig. 4*A*). By contrast, gametocyte production in *Pynsun1* knock-out line was not significantly reduced (Fig. 4*B*). The disruption of *Pynsun2* affected the production of male and female gametocytes similarly (Fig. 4*A*) while it showed no detectable effect on the propagation of asexual parasites (SI Appendix, Fig. S5*A*).

**Figure 4.**
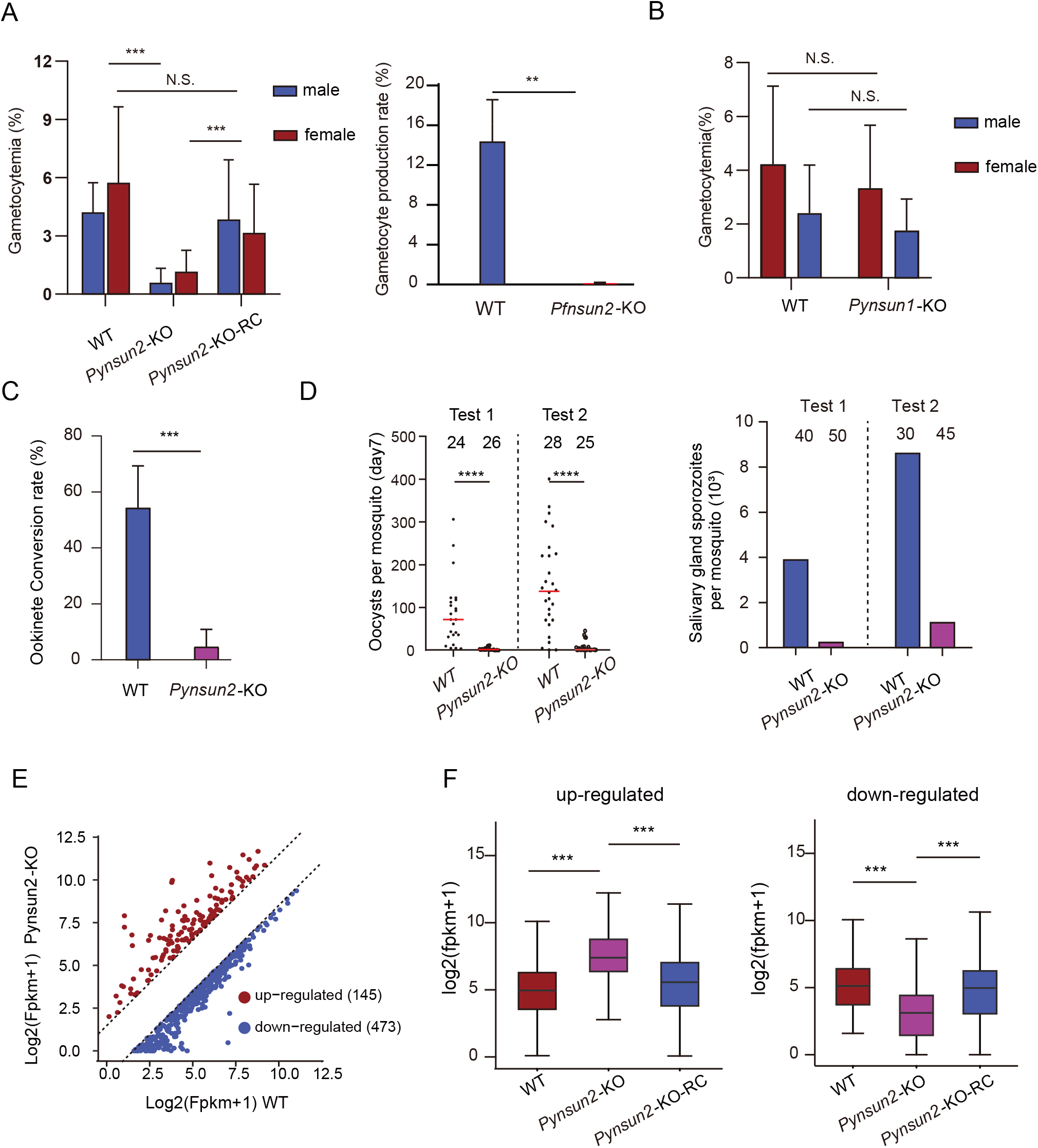
Loss of gametocytogenesis in *Plasmodium* NSUN2 knock-out lines and its restoration by gene complementation. **A**, Gametocytogenesis is markedly decreased in *Pynsun2*-KO and *Pfnsun2*-KO parasites relative to high levels in the original WT lines. Repair of the knock-out in *Pynsun2*-KO parasites restores gametocyte production. Error bars represent median and 95% CIs. (***: *P*-value < 0.001; **: *P*-value < 0.01; N.S.: no significant difference, Wilcoxon test.) **B**, No significant effect on gametocyte production detected after knock-out of the *Pynsun1* gene. Error bars represent median and 95% CIs. (N.S.: no significant difference, Wilcoxon test.) **C**, Markedly decreased ookinete conversion rates in the *Pynsun2*-KO relative to WT parasites. Data were obtained from two independent replicates. Error bars represent median and 95% CIs. (***: *P* < 0.001, Wilcoxon test). **D**, Number of oocysts in mosquito midguts 7 days after blood feeding with WT or *Pynsun2*-KO gametocytes (left), and number of sporozoites in mosquito salivary glands 14 days after blood feeding in the same experiments (right). The number of mosquitoes dissected in each group is indicated. Medians and 95% ICs are indicated. (***: *P* < 0.0001, Wilcoxon test). **E**, Identification of transcripts that are differentially expressed between *Pynsun2*-KO and WT parasites at the schizont stage. Transcripts that were found to have ≥ 3-fold up- or down-regulation are marked in red or blue, respectively. **F**, Upregulation and downregulation of transcripts in the knock-out clone are reversed by restoration of *Pynsun2* expression in the *Pynsun2-*KO-RC clone. (***: *P* < 0.001, Wilcoxon test).

Consistent with the effect of gene disruption on gametocyte production, *in vitro* ookinete conversion rates to mature forms were significantly reduced in the knock-outs relative to those of the WT parasites (Fig. 4*C*). Also, the numbers of midgut oocysts at day 7 and salivary gland sporozoites at day 14 were markedly decreased in *Pynsun2* knock-out parasites (Fig. 4*D*), suggesting that *Pynsun2* knock-out effects may impair the development of parasite stages in the mosquito. As with m^5^C modification of the mRNA transcripts, gametocyte production in the *Pynsun2-*KO-RC line was successfully rescued after restoration of *Pynsun2* expression (Fig. *4A*).

To further investigate the association of decreased numbers of gametocytes with diminished mRNA m^5^C modification, we identified transcripts whose m^5^C levels were significantly reduced in *Pynsun2-*KO relative to WT schizonts (SI Appendix, Table S7). Many of these genes showed significant *PyNsun2-*KO associated expression changes, including 145 instances of upregulation and 473 instances of downregulation (Fig. 4E), both of which were reversed after restoration of *Pynsun2* expression in the knock-out line (Fig. 4F). Among the downregulated genes, we identified transcripts that have been reported to be AP2-G-mediated and involved in gametocyte development (5) (SI Appendix, Fig. S5*B*). These included transcripts of Py17x_0313200 (DEAD/DEATH), disruption of which can greatly reduce the number of gametocyte parasites in *P. berghei* (5). Interestingly, no significant change of AP2-G expression itself was evident with the *Pynsun2* knock-out (SI Appendix, Fig. S5*C*).

## Discussion

The results of this study illuminate a landscape of m^5^C mRNA transcriptome methylations in the schizonts and gametocytes of two *Plasmodium* species, *P. yoelii* and *P. falciparum*, and show that these epitranscriptomic modifications are linked to development of gametocytes. Further, the searches for possible RNA methyltransferases and findings from experiments to knock out and repair the candidate genes indicate that a homolog of the NSUN2 family plays a major role in these m^5^C mRNA modifications. Although little or no effect on the growth of asexual blood stage parasites was detected in these experiments, markedly decreased gametocyte production was found after disruption of the *Pynsun2* homolog in *P. yoelii* (*Pynsun2-*KO) and disruption of the *Pfnsun2* homolog in *P. falciparum* (*Pfnsun2-*KO). Moreover, in evaluations of *Pynsun2-*KO infectivity, the conversion rate of ookinetes from immature to mature forms was also decreased, suggesting that m^5^C mRNA modifications may be important both to gametocyte production and to parasite development following zygote formation in the mosquito.

Dynamic m^5^C modifications are vital to gene regulatory networks in a diverse variety of biological processes including embryogenesis in zebrafish (21) and stages of development in *Arabidopsis* (13). Global transcriptome fluctuations of 30% in m^5^C methylation occur with altered photosynthesis efficiency and heat tolerance in rice (19); and m^5^C fluctuations have likewise been associated with the maternal-to-zygotic transition and the development of bladder carcinoma (18, 21). The m^5^C modification patterns observed in the present study, including increased levels of methylation in gametocytes and their loss following knock out of *Plasmodium* NSUN2, are also associated with the essential processes of male and female gametocytogenesis and progression to mature ookinetes following fertilization.

m^5^C modifications can stabilize mRNA transcripts and increase half-life, and they may also influence export of the transcripts for translation in the cytoplasm (14, 21). In the present work, several transcripts known to be involved in sexual commitment and gametocyte development were found to be m^5^C-modified, suggesting a role for this form of post-transcriptional methylation in the timing and control of their expression. Interestingly, no m^5^C modification was found for AP2-G, a master regulator of sexual commitment and development (32–34), nor was there any evidence for a change of AP2-G expression in NSUN2-disrupted parasites. These observations suggest that pathways downstream of AP2-G are epigenetically modulated at the post-transcriptional level. Knock-outs of NSUN2 and loss of m^5^C methylation may adversely affect the stability and translation of mRNA transcripts in these pathways, resulting in dysregulation and impairment of effective gametocytogenesis.

As in other organisms (13, 14, 21), the m^5^C sites of transcripts targeted by NSUN2 are found predominantly in the protein coding regions of *Plasmodium* mRNAs. However, the profiles of m^5^C distributions within these regions differ among organisms. In the two *Plasmodium* species studied here, a peak of m^5^C levels in CDSs occurs just upstream of the stop codon. In contrast, the m^5^C levels in cells from human and mouse cells or from zebrafish embryos are more uniformly distributed across the CDSs or are enriched near the translation initiation sites. *Arabidopsis* shows two peaks of m^5^C enrichment that flank the stop codon. Differences in the consensus site logos for m^5^C modification are also present between *Plasmodium* spp. and other eukaryotes (18, 19, 35). In *P. yoelii* and *P. falciparum*, the target C nucleotide is flanked on each side by a highly AU-rich sequence, which may in part reflect the correspondingly high AT content of the genomes and transcriptomes of these two species.

Our homology searches for sequences related to NSUN2 identified other potential methylases, termed *Plasmodium* NSUN1, NSUN3, and NSUN4 in this work. Multiple attempts with CRISPR/CAS9 strategies to knock out these different homologs in *P. yoelii* and *P. falciparum* yielded only one parasite line with a disruption of *Pynsun1*. However, unlike the *Pynsun2* and *Pfnsun2* knock-outs, the *Pynsun1-*KO line showed little or no reduction of gametocyte production relative to that of WT *P. yoelii.* Further investigations are needed to establish phenotypes associated with *Plasmodium* NSUN1 as well as NSUN3 and NSUN4, which, in light of the multiple unsuccessful knock out attempts, are likely to be essential for parasite survival. We note that the levels of m^5^C methylation in schizonts of the *Pynsun2-*KO and *Pfnsun2-*KO lines were decreased by only 30–50% relative to WT schizonts, as measured by both RNA-BisSeq and LC-MS/MS. Writer(s) in addition to NSUN2 may thus be involved in m^5^C transcript modifications of asexual as well as sexual stage parasites. These possibilities, in addition to the potential activities of m^5^C methyltransferases on other RNAs such as tRNA and ncRNAs (36) also remain to be explored.

We asked if m^5^C modifications of schizonts and gametocytes might be subject to an ‘eraser’ (demethylase) activity, but currently no such m^5^C demethylase has been identified in any eukaryote in which the landscape and biological roles of m^5^C mRNA modification have been well studied. While this does not rule out the possibility that erasers may yet be found, it is possible that m^5^C levels may simply rise and fall due to different rates of m^5^C writer activity and subsequent degradation of m^5^C-modified transcripts. This possibility is supported by the positive correlation between m^5^C status and mRNA stability (half-life) in gametocytes, as well as the greater m^5^C/C levels detected by LC-MS/MS in gametocytes relative to shorter-lived schizonts.

Control of mRNA turnover and translational repression has been shown to be essential for *Plasmodium* sexual development and is mediated by a member of the DDX6-family of DEAD box RNA helicases (37). Translational repression also occurs in mature sporozoites and is associated with their quiescence in mosquito salivary glands until blood feeding and salivation (38–40).

Here we have demonstrated that m^5^C modifications contribute to the stabilization of mRNA transcripts in *Plasmodium*, play an essential role in sexual stage development, and may be important for the maturation of ookinetes. We also speculate that m^5^C “reader” molecules may recognize these modifications and facilitate the transport of methylated transcripts to the cytoplasm for eventual translation. Further investigation of these m^5^C-mediated processes and those of other epitranscriptomic modifications such as m^6^A (12) could yield new insights into gametocytogenesis, mosquito infectivity, and molecular pathways that might be exploited to interrupt malaria transmission.

## Methods

### Animal use and ethics statement

All animal experiments were performed in accordance with approved protocols (XMULAC20140004) by the Committee for Care and Use of Laboratory Animals of Xiamen University. ICR mice (female, 5 to 6 weeks old) were purchased and housed in the Animal Care Center of Xiamen University and kept at room temperature under a 12 h light/dark cycle at a constant relative humidity of 45% and were used for parasite propagation, drug selection, parasite cloning, and mosquito feedings.

### Parasite culture and mouse infections

*P. falciparum* strain 3D7 (NF54) parasites were cultured and synchronized as previously reported (12). Human erythrocytes were obtained under approved clinical protocol at the Shanghai Blood Center. Microscopy of Giemsa-stained thin blood films was used to monitor parasite development.

*P. yoelii* 17XNL strain parasites were obtained from the Malaria Research and Reference Reagent Resource Center (MR4) (https://www.beiresources.org/About/MR4.aspx). The parasites were injected intravenously or intraperitoneally into mice for infection, and parasitemia was monitored by Giemsa-stained thin blood films.

### Dot blot detection of m^5^C Levels

Dot blot detection was performed as described previously (41). Biotin-labeled RNA oligonucleotides synthesized with or without m^5^C were used as positive and negative controls. mRNA was isolated from preparations of total RNA by the Dynabeads® mRNA Purification Kit (Thermo Fisher Scientific). mRNA was denatured at 95 °C in a heat block for three min and chilled on ice immediately to prevent the re-formation of secondary structures of mRNA. A sample containing 0.5µg mRNA was applied directly onto the Hybond-N+ membrane (Millipore) and secured to the membrane by two treatment rounds in a HL-2000 Hybrilinker^TM^ Hybridization Oven/UV Crosslinker (Analytik Jena US) operating in Autocrosslink mode (setting of 1,200 microjoules [x100]; 25-50 sec). After UV crosslinking, the membrane was incubated in 10 ml of 5% of non-fat milk in PBST (PBS with 0.05% Tween-20) for 1 h at room temperature with gentle shaking. The primary mouse anti-m^5^C antibody (1:1000; Abcam, ab10805) was then added in 10 ml of antibody dilution buffer (non-fat milk in PBST) and incubated overnight at 4 °C. After three times washing (5 min each) in 10 ml of PBST, membrane was finally incubated with HRP-conjugated anti-mouse IgG (1:5000; GE Healthcare) overnight at 4 °C. Signals from m^5^C were developed with an enhanced chemiluminescence (ECL) western blotting kit (GE Healthcare).

### mRNA modification determinations by LC-MS/MS

Total RNA was extracted from synchronized parasites samples using Trizol (ThermoFisher Scientific) followed by treatment with DNase I (ThermoFisher Scientific). mRNA was isolated from total RNA samples using NEBNext Poly(A) mRNA Magnetic Isolation Module (NEB). Purified mRNA was quantified using the Qubit RNA HS Assay kit (ThermoFisher Scientific). mRNA was hydrolysed to single nucleotides and the pretreated nucleoside solution was deproteinized using a Satorius 10,000-Da MWCO spin filter. Analysis of nucleoside mixtures was performed on an Agilent 6460 QQQ mass spectrometer with an Agilent 1260 HPLC system. Multi reaction monitoring (MRM) mode was performed due to its high selectivity and sensitivity attained through working with parent-to-product ion transitions. LC-MS data was acquired using Agilent Qualitative Analysis software. MRM peaks of each modified nucleoside were extracted and normalized to the amount of mRNA purified. Detailed analysis of mRNA modifications was performed as previously described (12, 35).

### Transcriptome sequencing

Total RNA was isolated from parasite preparations by the Direct-zol RNA Kit (Zymo Research). The poly(A) RNA fraction was selected using KAPA mRNA Capture Beads (KAPA), and fragmented to about 300–400 nucleotides (nt) in length. A KAPA Stranded mRNA-Seq Kit (KAPA) was used for RNA-seq library preparation for strand-specific RNA-seq. The libraries were sequenced on an Illumina NovaSeq 6000 platform to generate 150 bp pair-end reads.

### RNA-BisSeq data collection from transcripts and analysis

Parasites were isolated by treatment of 200 – 300 µl of loosely packed red blood cells from blood or culture at 3% parasitemia with 0.15% saponin in 1 × PBS on ice for 10 min. The parasite pellets were washed twice with precooling 1 × PBS and then resuspended in 1 mL TRIzol (Invitrogen). Total RNA was extracted using the Direct-zol RNA Kit (Zymo Research) according to the manufacturer’s instructions. mRNA enrichment from the total RNA was performed using the Dynabeads™ mRNA Purifcation Kit (Invitrogen) according to the manufacturer’s instructions.

RNA bisulfite treatment and purification of converted RNA was performed using the Methylamp™ RNA Bisulfite Conversion Kit (Epigentek Group Inc.) according to the manufacturer’s instructions. One μg of mRNAs along with 5 ng of dihydrofolate reductase (*dhfr*) RNA (1:2000) were used as an input RNA and unmethylated control, respectively. The resulting RNA was subsequently used for fragmentation and library construction using the KAPA Stranded mRNA-Seq Kit (KAPA) according to the instructions provided by the manufacturer. Sequencing was performed on an Illumina NovaSeq 6000 instrument with paired end 150-bp read length.

Raw sequencing data were filtered to remove low-quality reads and adapter contaminations with cutadapt (42) as detailed in Methods. Reads with average quality score ≥ 20 and length ≥ 50bp were kept. Clean reads were mapped to the *P. falciparum* 3D7 genome (Pf 3D7 v32, obtained from PlasmoDB) or to the *P. yoelii* 17X genome (Py17x v45, obtained from PlasmoDB) using meRanGh, and only unambiguously aligned reads were used to call m^5^C sites using the meRanCall tool from the meRanTK toolkit (43) (FDR < 0.01). Analysis of the *dhfr* spike-in revealed C to T conversions were around 98% (SI Appendix, Table S1). At least 20 reads were obtained for the candidate cytosine positions, and were recorded if methylated cytosine depth was greater than 5 and m^5^C level was greater than 0.1. mRNA-BisSeq data collection and analysis were performed in two replicate experiments. Detection of an m^5^C site in both replicates was required for its inclusion in further analysis. The m^5^C sites were mapped to the regions of introns, exons, 5’-UTRs, and 3’-UTRs as previously described (14). The meRanCall tool from meRanTK was used to localize m^5^C sites preferences with an input parameter of -sc 10. Logo plots were generated with R package ggseqlogo (44). PyNSUN2 targeted genes were defined as transcripts with m^5^C modifications in WT schizont parasites that are lost in *Pynsun2*-KO parasites.

### Plasmid construction and transfection

To disrupt the *PfNSUN2* opening reading frame (ORF) CDS in WT 3D7 and NF54 strains, we used CRSIPR/Cas9 to generate knock-out vectors. Briefly, the pL6 plasmid (45) was modified to contain a complementary sequence of sgRNA between the Xho I and Avr II sites. For each knock-out assay, at least two sgRNAs were individually incorporated into plasmids to target the CDS of their target gene (SI Appendix, Table S6). Sequences (∼ 1kb) homologous to the target genes were designed to include several base pair deletions and were inserted into the pL6 plasmid between Asc I and Afl II sites by the In-Fusion PCR Cloning System (primer pairs are listed in SI Appendix, Table S6). Recombinant plasmids with were simultaneously transfected along with pUF-Cas9 vectors into ring-stage parasites of 3D7-G7 and NF54-F5 by electroporation as described previously (46). Drug-resistant parasites were obtained after three or four weeks of continuous culture under 2.5 nM WR99210 pressure (12). Selected knock-out or knock-in parasites were confirmed by PCR detection and cloned by limiting dilution.

*P. yoelii* genetic modifications were performed using the CRISPR/Cas9 plasmid pYCm as previously described (47). To obtain gene knock-outs, 5’ and 3’-genomic fragments (400 to 700 bp) of target genes were amplified using the corresponding primers (SI Appendix, Table S6). The PCR products were restriction-digested and cloned into matched sites of the pYCm vector. SgRNA oligonucleotides were annealed and inserted into the pYCm vector. At least two sgRNAs were designed to disrupt the CDS of a target gene for each deletion modification using the online program EuPaGDT (48). For PyNSUN2 tagging, a 400 to 800 bp fragment from C-terminal of the CDS and 400 to 800 bp segment from the 3’-UTR were amplified and fused with a DNA sequence 6HA in frame at C-terminal of the gene. There were at least two sgRNAs designed to target sites close to the C-terminal of the gene CDS region for incorporation of the insert. Five micrograms of circular recombinant plasmid DNA were electroporated into infected RBCs using a Lonza Nucleofector, and the transfected parasites were intravenously injected into a naïve ICR mouse. Infected mice were treated with pyrimethamine (6 μg/ml) in drinking water; drug-resistant parasites were obtained following 5 to 7 days of drug selection.

### *In vitro* transcription of mouse *dhfr* RNA

Institute of Cancer Research (ICR) mouse heart tissue (0.05g) was obtained and then resuspended in 1 mL TRIzol (Invitrogen) for RNA extraction. The full-length *Mus musculus dhfr* CDS was amplified from cDNA and cloned between the XbaI and XhoI sites of the pBSKS vector (Stratagene, La Jolla, CA) containing a T7 promoter (forward primer: 5’-CACCGCGGTGGCGGCCGCTCTAGAATGGTTC GACCATTGAACTGC-3’; reverse primer: 5’-GTACCGGGCCCCCCCTCGAGTTAGTCTTTCTTCTCGTAGACTTC-3’). The pBSKS-dhfr vector was linearized by FastDigest XhoI (Thermo Scientific™). Five hundred nanograms of linearized pBSKS-dhfr vector was used as a DNA template for *in vitro* transcription with T7 RNA Polymerase (New England Biolabs) at 37 °C for 4 h in a 20 μl reaction mixture, according to the manufacturer’s instructions. The DNA template was removed by DNase I (Invitrogen™) digestion for 15 min at room temperature.

### Western blotting

Protein fractionation was performed with modifications according to Till *et al* (49) and Baumgarten *et al* (12). Synchronous middle-late stage parasites were released from 1 mL infected red blood cells by 0.15% saponin lysis and washed twice in 1 × pre-cooled PBS. The parasite pellet was first resuspended in 0.5 mL cytoplasmic lysis buffer (20mM Hepes pH7.9, 10mM KCl, 1.5mM MgCl2, 1mM EDTA, 1mM EGTA, 0.65% NP-40, 1mM DTT, 1x Protease inhibitor cocktail) and incubated for 30 min on ice. Parasites were further homogenized for 100 strokes in a glass douncer, and the supernatant containing cytoplasmic proteins was isolated via centrifugation (10,000 rpm, 10 min, 4 °C). Protein samples were separated on SDS-PAGE and transferred onto PVDF membrane and visualized using ECL Western Blotting Detection Kit (GE Healthcare). The primary antibodies used in this study were mouse anti-HA (1:2000, Roche) and anti-GAPDH (1:1000, Servicebio, cat# GB12002), and secondary antibody was HRP-conjugated goat anti-mouse IgG (1:5000, Abcam, cat# ab6789).

### Gametocyte induction

For the *P. yoelii 17XNL* strain, gametocyte induction in mice was performed as previously described (47). Briefly, phenylhydrazine (80 μg/g mouse body weight) was administered to the ICR mice through intraperitoneal injection. The mice were infected three days later with 3.0 × 10^6^ parasites through tail vein injection. Gametocytemia typically peaked at day 3 post injection. Giemsa staining of blood smears was used to count the number of male and female gametocytes. Gametocytemia was defined as the ratio of sex-specific gametocyte (male or female) over total infected erythrocytes. All experiments were repeated three times independently.

For *P. falciparum,* gametocyte induction was performed as described previously with minor modifications (50, 51). The *P. falciparum* population was synchronized and expanded to 8% parasitemia in culture (4% haematocrit). Medium changes were performed daily without dilution of the culture and without disturbing the red blood cell layer at the bottom of the culture dish. Giemsa’s solution staining of thin blood smears was used to count the number of gametocytes from day 3. Stage III gametocytes were harvested on days 5–6, stage IV on days 8–9, and stage V on days 12–13. Gametocytemia determined on day 8-9 was used to calculate sexual commitment (%). Each experiment had three biological replicates.

### Ookinete conversion rate determinations

Conversion rates to mature ookinetes were determined as recently described (52). Samples taken from culture 12 h after fertilization were stained by Giemsa solution (Sigma, cat# GS80). Each conversion rate was calculated as the number of mature ookinetes (stage V) over the number of total ookinetes (stages I–V).

### RNA-Seq for mRNA half-life

ICR mouse blood with 4–6% gametocytemia was collected in anticoagulant tubes with heparin and immediately added to gametocyte culture medium (RPMI 1640 medium) in a blood/medium volume ratio of 1:10. Actinomycin D (Sigma-Aldrich) was added at 20 μg/ml (53), and samples were collected 0, 1, 3, and 5 h later. Total RNA was extracted by Direct-zol RNA Kit (Zymo Research) as per the manufacturer’s instructions. Prior to construction of the library with KAPA Stranded mRNA-Seq Kit Illumina platform (KK8421), ERCC RNA Spike-In Control Mixes (Ambion) was added proportionally to the total RNA of each sample (0.1 µL per sample) (54). Two biological replicates were performed for gametocyte samples.

### Mosquito maintenance and mosquito feeding

Sucrose solution (10%) was used to feed *Anopheles stephensi* mosquitoes (strain Hor) (55), which were reared at 28 °C, 80% relative humidity, in a 12 h light/dark cycle and blood feeding assay was performed at 23°C. Each anaesthetized mouse with 4–6% gametocytemia was used to feed 80 female mosquitoes for 30 min. For oocyst detection, mosquito midguts were dissected seven days after blood-feeding and stained using 0.1% mercurochrome. For salivary gland sporozoite counting, salivary glands from 30 – 50 mosquitoes were dissected on day 14 post blood-feeding, and the number of sporozoites per mosquito was calculated.

### RNA sequence analysis

Low-quality and adaptor sequences were trimmed using cutadapt (v1.18) (42) with parameters: -a AGATCGGAAGAGC -AAGATCGGAAGAGC –trim-n -m 50 -q 20,20. RNA sequencing reads were aligned using hisat2 (v2.1.0) (56) with strand specific mode (--rna-strandedness RF) to the *P. falciparum* 3D7 genome (Pf 3D7 v32, obtained from PlasmoDB) or to the *P. yoelii* 17X genome (Py17x v45, obtained from PlasmoDB). Mapped reads were subsequently assembled into transcripts guided by the PlasmoDB gff annotation files (Pf3D7 v32/ Py17x v45) using featureCounts (v1.6.1) (57) with parameters: -M -p –B -C -s 2. Read counts were obtained using featureCounts (v1.6.1). FPKM for genes were calculated according to the formula 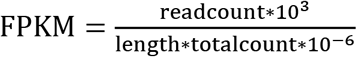. Genes with foldchange ≥ 3 were considered as differentially expressed genes for compared groups.

### Gene homolog identification and phylogenetic analysis of the m^5^C writer

Candidate proteins of the malaria parasite m^5^C writer complex were identified by blast search against PlasmoDB database (https://plasmodb.org/plasmo/) using the amino acid sequence of human mRNA m^5^C methylase NSUN2 as a query. The amino acid sequences of human, rice, and *Arabidopsis* RNA (cytosine-5)-methyltransferases (RCMTs) domain family members were obtained from the NCBI database (https://www.ncbi.nlm.nih.gov/). Clustal X software (58) was used to construct the Neighbor-joining phylogenetic tree with 1,000 bootstrap replicates.

### mRNA half-life analysis

Raw sequencing data were filtered to remove low-quality reads and adapter contaminations with cutadapt (v1.18) (42), as described above. Reads with average quality score ≥ 20 and length ≥ 50 bp were kept. Clean reads were mapped to the *P. yoelii* 17X genome (Py17x v45, obtained from PlasmoDB) with ERCC spike in sequence using hisat2 (v2.1.0) with strand specific mode (--rna-strandness RF). Uniquely mapped reads were used to calculate read count for each transcript using software featureCounts. To remove low abundance transcripts, transcripts with at least five read counts at 0 h were kept for downstream half-life analysis. The read count of each transcript in a time point was normalized with the sum of reads mapping to all ERCC spike-in transcripts. Half-lives of transcripts were calculated using R package MINPACK (59). The decay rates for transcripts were constant upon transcription inhibition. Transcripts are assumed to exponentially decay. Therefore, the transcripts remaining at specific time point were calculated by the equation: 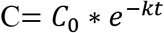. Here, C represents transcript abundance as a function of time, C_0_ is the transcript abundance at 0 h, and k is the first order rate constant, from which half-life values can be derived with the equation: 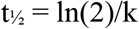.

### Proteomics analysis

TMT-based quantitative proteomics analysis was performed by Novogene Bioinformatics Technology Co Ltd. (Beijing, China), as described previously (60). Briefly, gametocytes of *P. yoelii* were collected and ground individually with liquid nitrogen and lysed in protein lysis buffer (100 mM NH_4_HCO_3_, 8 M Urea and 0.2% SDS, pH = 8.0) followed by ultrasonication on ice for 5 min. Desalted peptides were labeled with TMT 6-plex reagents (TMT6plex™ Isobaric Label Reagent Set, Thermo Fisher) according to the manufacturer’s instructions. Labeled peptides of all samples were mixed equally and desalted by peptide desalting spin columns (Thermo Fisher, 89,852). A C18 column (Waters BEH C18, 4.6 × 250 mm, 5 μm) on a Rigol L3000 HPLC was used to fractionate for TMT-labeled peptide mix. An EASY-nLC^TM^ 1200 UHPLC system (Thermo Fisher) coupled with an Orbitrap Q Exactive^TM^ HF-X mass spectrometer (Thermo Fisher) operated in the data-dependent acquisition (DDA) mode was used for shotgun proteomics analyses. The resulting spectra from each fraction were searched separately against *P. yoelii* 17X proteome (Py17x v45, obtained from PlasmoDB) by the search engines: Proteome Discoverer 2.2 (PD 2.2, thermo). At least one unique peptide with false discovery rates (FDR) < 1.0 % on the peptide and protein level, respectively, was considered as a confident peptide spectrum match for further analysis. The intensities of the TMT reporter ions against those ones provided in the databases were compared for protein quantification.

### Gene Ontology (GO) analysis

GO analysis of specific genes were performed using EnrichGO tool from Plasmodb (https://plasmodb.org/). Only terms in the biological process category were shown. GO terms with p-value ≤ 0.05 were considered as statistically significant terms. Genes enriched for terms sexual development (GO: GNF0004) were obtained from a previous study (61) and used for enrichment analysis..

## Supporting information

Stables and figures

## Acknowledgments

This work was supported by the National Natural Science Foundation of China (NSFC) [81630063, 81971959, 31671353] to Q.Z., the Division of Intramural Research at the U.S. National Institute of Allergy and Infectious Diseases (J.M. and T.E.W.), and National Key R&D Program of China Grants [2018YFA0507300] to Q.Z., NSFC [31771419, 31721003] to C.J., NSFC [31772443, 31970387] to J.Y.

## Author Contributions

QZ, CJ, JY, JM and TW conceived and designed experiments. GG, PQ, BL and XH performed the majority of experiments with the help from XS, QM, XY and YZ. ML, GY, SS, WL, LW, LG and JM conducted most bioinformatics analysis. QZ, CJ, JY, JM and TW wrote the manuscript with contribution from ML, GG, LJ, JC and RC. All authors discussed and approved the manuscript.

## Codes and Data Availability

All data is available in the main text or the supplementary materials. The raw and processed high-throughput sequencing data of this study have been deposited into the Gene Expression Omnibus (GEO) and are publicly accessible under accession number GSE159127 (token: gjcrewgkdnijrsh).

---

**Figure S1. Detection and transcriptome-wide mapping of m^5^C mRNA modifications in *Plasmodium*. A**, Venn diagrams display the numbers of mRNA m^5^C sites that are shared or unique in two independent replicate determinations (rep1, rep2) from schizont and gametocyte stages of *P. yoelii* (Py.S, Py.G) and *P. falciparum* (Pf.S, Pf.GIII, Pf.GIV, Pf.GV). Box plots indicate m^5^C levels at the shared and unique sites in each replicate pair. (***: p-value < 0.001, Wilcoxon test). Maximum, minimum, and median values are indicated in the box plots. **B,** Histograms and box plots indicate the distributions of modification levels at m^5^C sites in the schizont and gametocyte stages of *P. yoelii* and *P. falciparum*. **C,** Frequency histograms indicate the number of transcripts by the m^5^C sites they carry in schizonts and gametocytes of *P. yoelii* (Py.S, Py.G) and *P. falciparum* (Pf.S, Pf.GIII, Pf.GIV, Pf.GV). **D**, Proportions of m^5^C sites in each parasite stage are displayed by sequence context: CHH (orange), CHG (red) and CGH (green), where H represents A, C or U. **E**, Transcriptome-wide distributions of mRNA m^5^C sites in the predicted CDS, introns, and flanking regions of genes expressed in schizonts and gametocytes of *P. yoelii* (Py.S, Py.G) and *P. falciparum* (Pf.S, Pf.GIII, Pf.GIV, Pf.GV). The presence of m^5^C in predicted intron regions suggests the methylation of incompletely or alternatively spliced transcripts.

**Figure S2. Heatmaps representing m^5^C modifications in schizont and gametocyte stages of *P. yoelii* (Py.S, Py.G) and *P. falciparum* (Pf.S, Pf.GIII, Pf.GIV, Pf.GV).** Vertical bars mark regions with sequences that match with GO terms for sexual development and nucleic acid metabolic processes.

**Figure S3. Comparatively high m^5^C methylation levels correlate with expression of gametocytogenesis-associated genes. A**, Heat map displays are aligned to indicate transcription levels alongside the relative m^5^C levels in *P. yoelii* and *P. falciparum* schizonts and gametocytes. Data on transcript sequences and m^5^C levels were each obtained from two biological replicates using RNA-Seq and RNA-BisSeq, respectively. **B**, Abundance of transcripts in gametocytes relative to schizonts correlates with their stage-specific m^5^C methylation levels. Levels of individual transcripts in gametocytes of *P. yoelii* (G) or *P. falciparum* (stage V, GV) relative to their levels in schizonts (S) are plotted by log_2_ ratio. Results from transcripts with lesser or greater m^5^C levels in gametocytes than schizonts are shown in blue (S > G) and red (S < G), respectively, while those with little or no difference of m^5^C levels are shown in yellow (S = G).

**Figure S4. Disruption of the *Pfnsun2* gene in *P. falciparum* and restoration of *Pynsun2* expression in *P. yoelii* knock-out parasites. A**, Predicted protein sequences and electropherograms from the *P. falciparum* WT and *Pfnsun2-*KO parasites. An early stop codon is introduced by the genetic modification. Red arrow indicates the deletion site. **B**, Immunoblot detection of 6×HA tagged PyNSUN2 protein in the asexual blood stages of *Pynsun2*-KO-RC and in a separate parasite strain (*Pynsun2-*6HA) containing a 6×HA tagged version of endogenous PyNSUN2. Control bands of GAPDH detection verify comparable loading of the lanes.

**Figure S5. PyNSUN2 deficiency leads to a reduction in gametocyte production. A**, Results demonstrating comparable development of parasitemias in WT- and *Pynsun2*-KO-infected mice. Data points are means ± SEM from three independent biological experiments. **B**, Decreased expression of AP2-G induced genes associates with dramatically reduced m^5^C levels in the *Pynsun2-*KO (KO) parasites. Error bars represent SEM for two biological replicates. **C**, Histogram of AP2-G (PY17X_1440000) transcript levels in WT and *Pynsun2*-KO parasites. Medians and 95% ICs are indicated. (N.S.: no significant difference, Wilcoxon test.)

**SI Appendix, Table S1.** Summary of RNA-BisSeq mapping results and statistics of m^5^C sites per transcript from schizonts and gametocytes of *P. yoelii* and *P. falciparum*

**SI Appendix, Table S2.** m^5^C levels of transcripts in schizont and gametocyte stages of *P. falciparum* and *P. yoelii* (WT, *Pynsun2-*KO, and *Pynsun2-*KO-RC lines)

**SI Appendix, Table S3.** Transcripts with greater than 3-fold upregulation in gametocytes relative to schizonts of the *P. falciparum* (NF54) and *P. yoelii* (17XNL) lines

**SI Appendix, Table S4.** Individual half-lives of transcripts in *P. yoelii* gametocyte stages

**SI Appendix, Table S5.** Protein abundances in schizonts and gametocytes of *P. yoelii* (two replicates, TMT quantification)

**SI Appendix, Table S6,** Primer and gRNA sequences used for plasmid construction and gene knock-out experiments

**SI Appendix, Table S7.** Genes exhibiting ≥ 3-fold expression changes between the *P. yoelii* WT, *Pynsun2-*KO, and *Pynsun2-*KO-RC parasites.

