## Supplementary figures and images for "5-methylcytosine modification by *Plasmodium* NSUN2 stabilizes mRNA and mediates the development of gametocytes"

### MT2_project_figures_supp_py.pdf

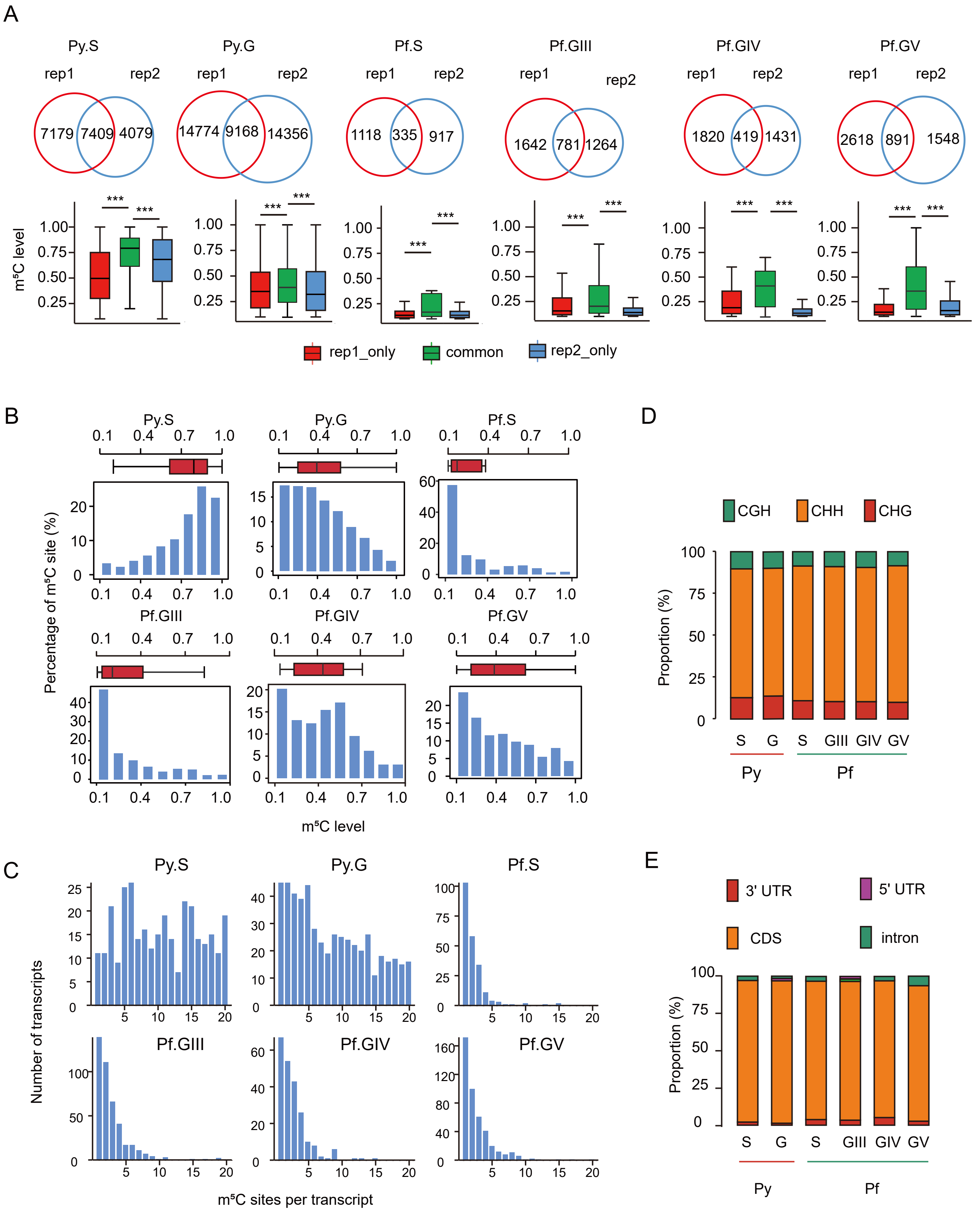

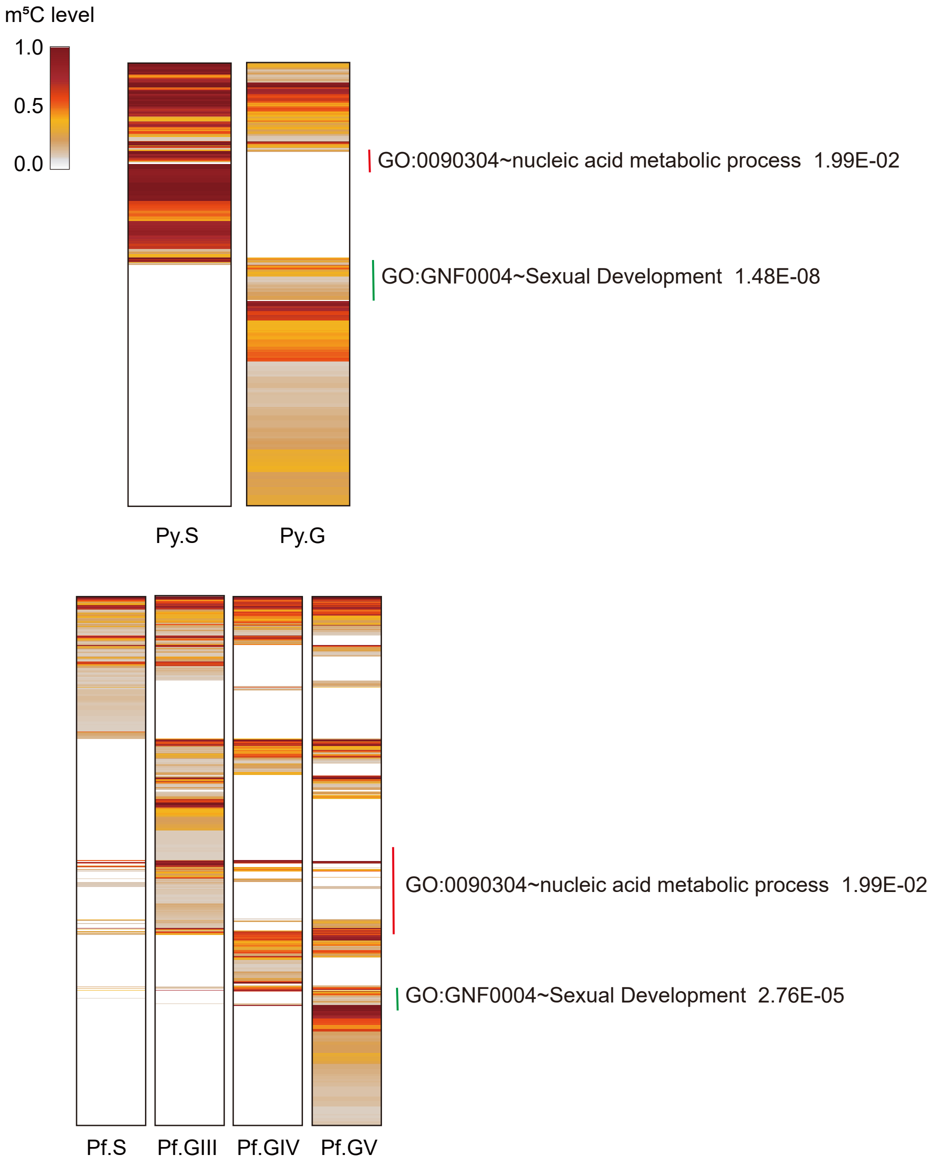

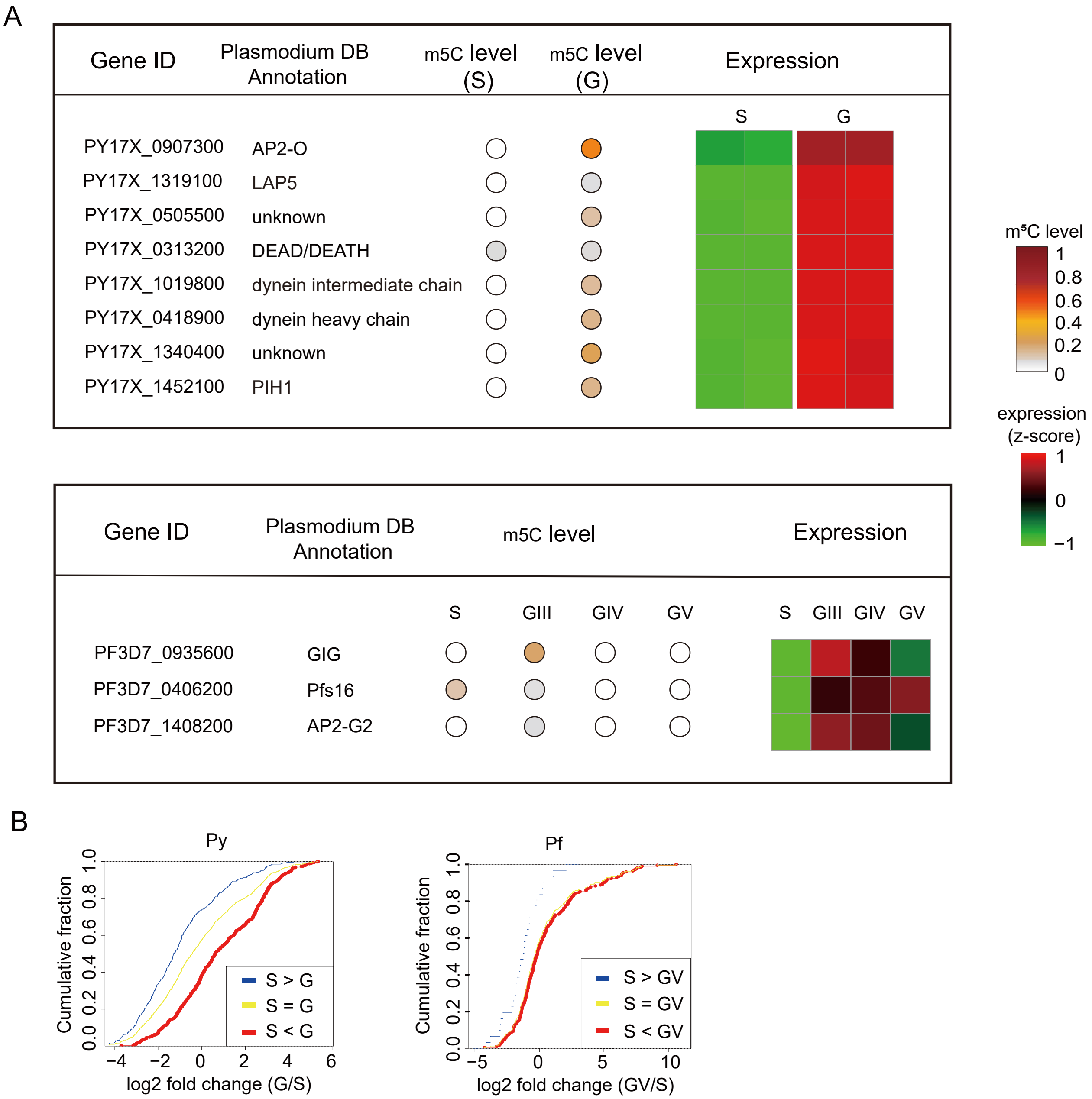

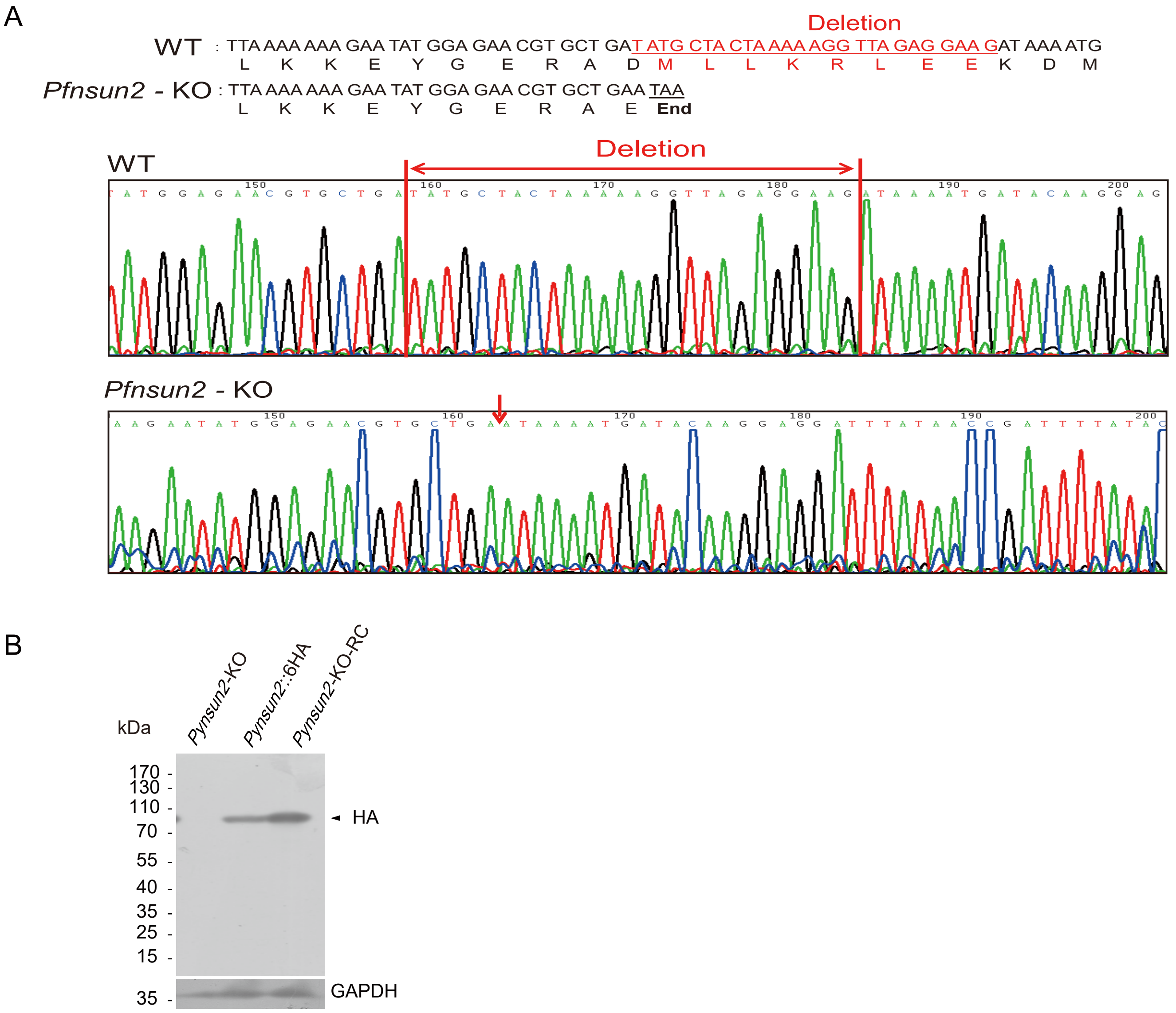

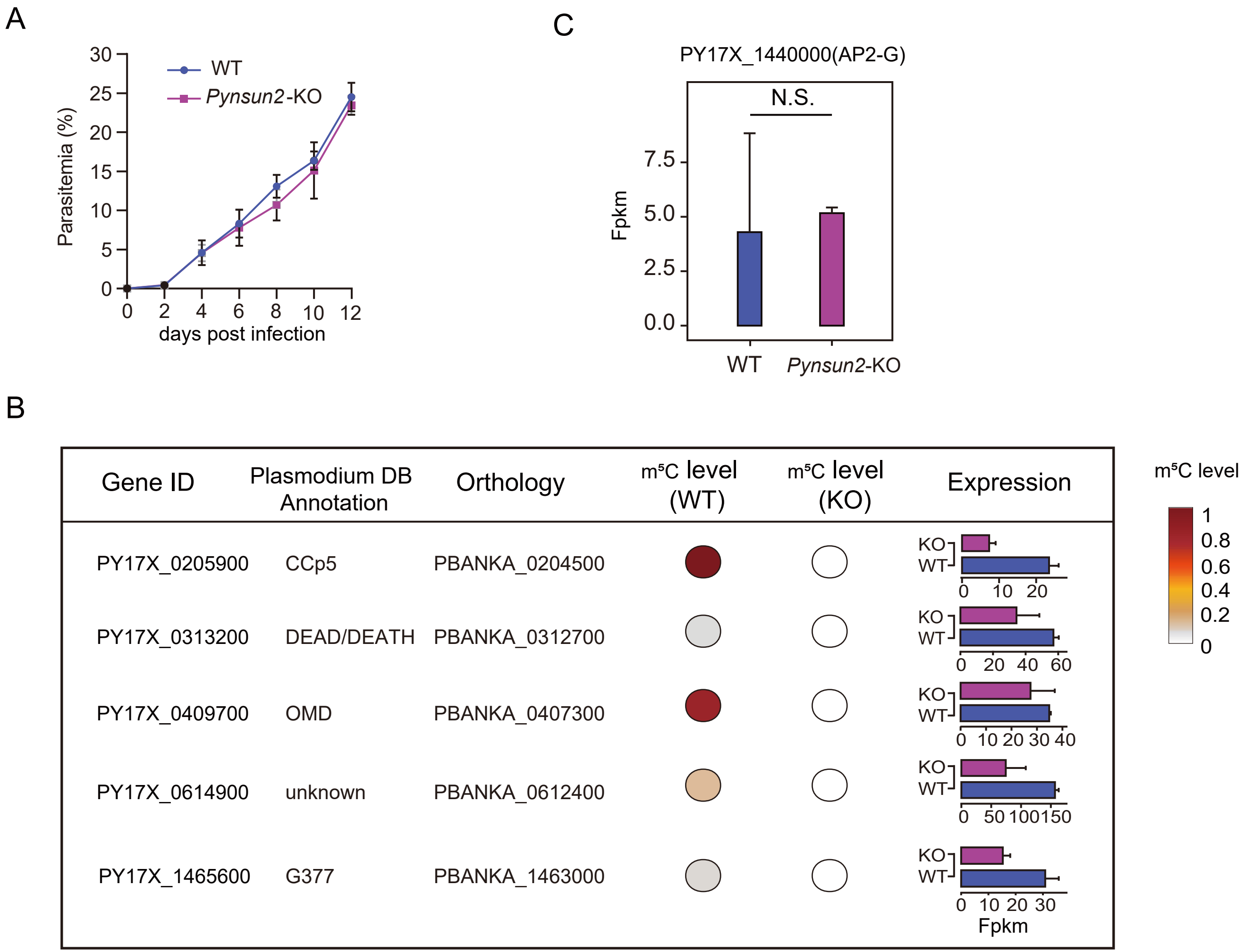
